# An agent-based framework for improving wildlife disease surveillance: A case study of chronic wasting disease in Missouri white-tailed deer

**DOI:** 10.1101/478610

**Authors:** Aniruddha V. Belsare, Matthew E. Gompper, Barbara Keller, Jason Sumners, Lonnie Hansen, Joshua J. Millspaugh

**Affiliations:** School of Natural Resources, University of Missouri, Columbia, Missouri, United States of America; Minnesota Department of Natural Resources, St. Paul, Minnesota, United States of America; Missouri Department of Conservation, 3500 East Gans Road, Columbia, Missouri, United States of America; W.A. Franke College of Forestry and Conservation, University of Montana, Missoula, Montana, United States of America

**Keywords:** agent-based models, chronic wasting disease, sample size, surveillance, white-tailed deer, wildlife diseases

## Abstract

Epidemiological surveillance for important wildlife diseases often relies on samples obtained from hunter-harvested animals. A problem, however, is that although convenient and cost-effective, hunter-harvest samples are not representative of the population due to heterogeneities in disease distribution and biased sampling. We developed an agent-based modeling framework that i) simulates a deer population in a user-generated landscape, and ii) uses a snapshot of the *in silico* deer population to simulate disease prevalence and distribution, harvest effort and sampling as per user-specified parameters. This framework can incorporate real-world heterogeneities in disease distribution, hunter harvest and harvest-based sampling, and therefore can be useful in informing wildlife disease surveillance strategies, specifically to determine population-specific sample sizes necessary for prompt detection of disease. Application of this framework is illustrated using the example of chronic wasting disease (CWD) surveillance in Missouri’s white-tailed deer (*Odocoileus virginianus*) population. We show how confidence in detecting CWD is grossly overestimated under the unrealistic, but standard, assumptions that sampling effort and disease are randomly and independently distributed. We then provide adjusted sample size recommendations based on more realistic assumptions. These models can be readily adapted to other regions as well as other wildlife disease systems.

## 1. Introduction

Surveillance of wildlife populations is an essential tool for effective management of diseases that pose conservation, economic or public health threats. Samples for wildlife disease surveillance are frequently obtained from hunter-harvested animals (e.g. [1–4]). This approach is practical and cost-effective as samples are obtained from a large geographic area and sampling costs are dramatically reduced. In recent decades, wildlife agencies across USA and Canada have implemented harvest-based chronic wasting disease (CWD) surveillance in cervid populations [5, 6]. CWD is an emerging prion disease of cervids, including white-tailed deer (*Odocoileus virginianus*), mule deer (*O. hemionus*) and elk (*Cervus elaphus*). Currently reported from 24 states in the continental United States and two provinces in Canada, the continuing spread of CWD is a serious long-term threat to the health of free-ranging cervid populations [7–10].

For emerging wildlife diseases like CWD, surveillance programs are aimed at prompt detection in populations where the disease has not been previously documented. Epidemiological data generated from sampling the population of interest is used to infer disease status of the population (presence/absence, prevalence, distribution). Reliable inferences can be made only if sufficient representative samples are obtained. Sample sizes for epidemiological surveillance are often determined from standardized tables (or online programs) that use hypergeometric distribution (sampling without replacement) for finite populations or a binomial distribution (sampling with replacement) for large populations (>100000), under the assumption that samples are representative of the population of interest [5,11,12].

However, hunter-harvested samples will rarely be representative of the population and may result in biased estimates and erroneous inferences about disease presence and distribution [6, 13]. Biases in hunter-harvested samples can arise from disease distribution heterogeneities in the population (infected individuals are often clustered rather than being uniformly or randomly distributed through a population) and factors like harvest regulations, hunter selectivity, variable land access, and uneven hunter distribution. Wildlife agencies need to consider such biases and adapt their disease surveillance strategies accordingly. This issue is particularly important when the disease prevalence is very low and yet the potential for impact on wildlife, domestic animals, or public health is high.

The white-tailed deer is one of Missouri’s most valuable natural resources, and the continuing spread of CWD poses a serious threat to the health of Missouri’s free-ranging deer populations. Logistical and financial constraints limit the ability of mangers to implement strict probability-based sampling schemes, hence CWD surveillance continues to rely on hunter-harvested samples. Statewide surveillance of hunter-harvested deer for CWD was initiated by the Missouri Department of Conservation in 2002 [14], but the first case of CWD in free-ranging deer was detected in 2012. As of December 2017, more than 76,600 free-ranging deer have been tested and 43 found positive for CWD in Missouri (0.06% of sampled deer) [15–17]. In 2016-17 alone, the state tested >25,600 deer and detected nine positive cases in five counties (0.04% of sampled deer). These numbers illustrate both the daunting challenge involved with surveillance, and the level of concern over the disease in the state.

CWD occurs at a low prevalence in most areas, and has a clustered distribution with focal areas of infection near the point of introduction [18–20]. Three considerations are therefore vital in the context of CWD surveillance: (1) the very low prevalence of disease when detection matters most, (2) the non-random distribution of diseased individuals, and (3) the non-random nature of samples derived from hunter harvested animals. Agent-based models (ABMs, also called Individual-based models) are well-suited to solving these problems. ABMs are formulated using relevant information about individuals, their behavior and the landscape they occupy, and therefore can readily incorporate non-random and heterogeneous aspects of the population, the disease, and harvest. Moreover, ABMs have been used to inform policy in many fields including natural resource management, epidemiology and public health.

We have developed an agent-based modeling framework that incorporates real-world heterogeneities arising due to disease distribution on the landscape and sampling biases in hunter-harvest, and provides a rigorous, quantitative framework for informing disease surveillance strategies. Our ABM framework has two constituent models: (1) MO*Ov*POP (MissOuri *Odocoileus virginianus* POPulation simulation) simulates a realistic deer population for the region of interest and (2) MO*Ov*POP*surveillance* simulates disease coverage in the model deer population and then samples the population for disease through hunter-harvest. Combining these two models allows managers to iteratively estimate population-specific sample size for detection of disease with a high-level of confidence. Here, we illustrate the use of this framework to derive optimum sample sizes for efficient and reliable CWD surveillance in Missouri’s white-tailed deer populations. Though we focus on CWD in white-tailed deer populations of Missouri, this model-based framework can be readily adapted to other regions, and other host-pathogen systems.

## 2. Background information and modeling considerations

The purpose of the first model, MO*Ov*POP, is to generate a snapshot of a target deer population that is as realistic as possible. The model landscape is a grid of one square mile patches derived from the GIS coverage data (forest cover) for the area of interest. In the Midwest North America, deer thrive well in landscapes where forest cover and agricultural food are juxtaposed, but less so in heavily forested or predominantly agricultural areas [21, 22]. Therefore, contiguous patches in the model landscape with average forest cover between 25% and 75% are designated as deer habitat.

The model is initialized using demographic attributes derived from field data, specifically, density, sex-ratio, age composition, reproduction and mortality (both hunting and non-hunting). Deer in the model are treated as individual agents that occur on habitat patches; they interact with each other and the environment. Deer are classified in four age-classes: young fawns (up to 6 months old), older fawns (7 to 12 months old), yearlings (13 to 24 months old) and adults (25 months or older). Social organization, group dynamics and dispersal are explicitly incorporated in the model as these processes influence interactions among individual deer and underpin the heterogeneity in the population. These heterogeneities should be accounted for when designing surveillance for a disease like CWD.

White-tailed deer are social animals exhibiting an intermediate level of sociality, typically occurring in small, relatively stable groups of adult females and their recent offspring (doe social groups), loose bachelor groups of adult males, or as solitary individuals (male and female) [23, 24]. Furthermore, the pattern and strength of social affiliations in white-tailed deer populations fluctuate temporally. For instance, pregnant females seek isolation during the fawning season and become aggressive towards other deer including group members [25, 26]. Similarly, bachelor groups break up and males are solitary during the breeding season [27]. Pre-breeding interactions (sparring matches, flehmen behavior or tending behavior) and mating also influence the pattern of interactions between deer. We have accounted for such temporal fluctuations that influence group dynamics in this model.

Dispersal is an important component of white-tailed deer ecology, and can create opportunities for the spread of pathogens like CWD [28]. Though dispersal behavior is predominantly exhibited by yearling bucks [29, 30], female dispersal also occurs, albeit at lower rates [31]. Yearling buck dispersal can occur during the parturition period, cued by the intersexual aggression of pregnant females, or during the rut period, cued by the intrasexual aggression by adult males [32]. Agonistic behavior by pregnant does just before parturition is thought to be the reason for yearling female dispersal [33]. Every year of the model run, yearling buck and yearling female dispersal is scheduled before parturition in the fifth month. Additionally, yearling buck dispersal also occurs before rutting activity in the 11^th^ month every year. Dispersal rates and distances for yearling bucks and yearling females are extrapolated from published literature (details provided in Parameterization and Calibration section).

Similarly, parameters influencing deer reproduction and harvest have been extrapolated from the multi-year harvest data collected by MDC. We simulate the population forward in time until it achieves an equilibrium where the age-sex composition and population growth are all stable. A snapshot of the model-generated deer population is then exported and stored for subsequent use by the surveillance model.

The second model, MO*Ov*POP*surveillance*, imports MO*Ov*POP-generated deer population snapshot, simulates disease coverage (distribution of diseased individuals across age-sex classes and across the model landscape) and then implements harvest-based sampling (random or non-random) for detection of the disease.

MO*Ov*POP*surveillance* can simulate a range of CWD prevalence and distribution scenarios. In particular, low prevalence scenarios (∼populations where CWD has been recently introduced) are important in the context of CWD surveillance. The incorporation of non-random and heterogeneous real-world processes in the model facilitates sample size calculations for prompt and reliable detection of CWD.

## 3. Model description

MO*Ov*POP and MO*Ov*POP*surveillance* were both developed in NetLogo 6.0, a software platform for implementing agent-based models [34]. Model description is provided following the ODD (Overview, Design concepts, Details) protocol for individual-based models [35, 36]. Model codes are available via website repository “Open ABM CoMSES Computational Model Library” [37, 38].

### 3.1 Purpose

The purpose of the model-based framework is to provide a decision-making context for collection and analysis of harvest based epidemiological surveillance data. The framework is adapted for chronic wasting disease (CWD) surveillance of Missouri white-tailed deer populations.

### 3.2 State variables and scales

#### Entities

MO*Ov*POP and MO*Ov*POP*surveillance* have two entities: patches and deer. Deer are modeled as individuals that occur on patches in the model landscape.

#### State variables

Patches located on the edges of the model landscape are designated as border patches (*border*). Each patch is characterized by its percent forest cover (*forest-percent*) and the mean forest-percent (*dfp*). The mean forest-percent is required in the male yearling dispersal submodel and is calculated by averaging forest cover of a patch and its immediate neighbors. During the setup, patches are assessed for deer occupancy (*do*) based on their percent forest cover. Patch variable *dh* (deer habitat) is then used for grouping of contiguous deer occupancy patches, thus facilitating initial distribution of deer in the model landscape.

Each patch in MO*Ov*POP*surveillance* has two additional patch variables: *add* (average deer density) takes the value of average number of deer occurring on the patch and its immediate neighbors (8 patches), and *hp* designates the relative harvest pressure for a patch.

Each deer has eight state variables, which define individual characteristics that include age, sex, group membership and status. Deer in MO*Ov*POP*surveillance* have three additional state variables that designate CWD infection status (*cwd*), harvest status (*marked-for-harvest*?) and sampling status (*marked-for-testing?*).

Patch and deer state variables are listed in Table 1.

**Table 1.** State variables of agents included in MO*Ov*POP and MO*Ov*POP*surveillance*. All state variables except the deer state variables ‘aim’ and ‘add’ are unitless.

| Agent | Variable | Description |
| --- | --- | --- |
| Patch | forest-percent | forest cover expressed as a proportion |
|  | border | patches on the edge of the model landscape have border = 1, other patches have border = 0 |
|  | dfp | mean forest-percent for a patch and its immediate neighbors |
|  | do | deer occupancy; 1 if deer occur on a patch, 0 otherwise |
| | dh | deer habitat; $\geq 1$ if a patch or a group of contiguous patches qualifies as deer habitat, otherwise $< 1$ |
|  | add | average deer density calculated for a patch and its immediate neighbors (8 patches) (per square mile) |
|  | hp | designates harvest pressure on the patch: 0 no harvest, 1 low harvest and 2 high harvest |
| Deer | sex | 1 if male, 2 if female |
|  | aim | age in months |
|  | momid | dam's id number |
|  | gl | 1 if a doe social group leader, 0 otherwise |
|  | ml | 1 if a bachelor group leader, 0 otherwise |
| | groid | $\geq 0$ if member of a doe social group, -1 if a solitary female, -1 if a male yearling or adult deer |
| | gr | $\geq 1$ if a group leader, -1 if member of a doe social group, otherwise -2 |
|  | mgroid | 0 if a female deer, -2 if a male fawn, group leader id if member of a bachelor group, -1 for adult solitary males |
|  | cwd | CWD infection status of a deer: 0 uninfected and 1 infected |
|  | marked-for-harvest? | deer marked for harvest |
|  | marked-for-testing? | deer marked for sampling (CWD testing) |

#### Spatial scales

MO*Ov*POP has a spatially-explicit landscape that can be set up for individual counties or CWD management zones using GIS coverage data (forest cover) for the selected region. Irrespective of the region selected for simulation, each patch in the model landscape represents one square mile. We have used miles rather than kilometers as the measurement unit because of the past and current norms of the region and its management agencies, and the related need to make the results immediately applicable to those same agencies.

#### Temporal scale

MO*Ov*POP has a monthly time step, and the deer population is simulated for a period of 25 years before pre-harvest and post-harvest population snapshots are exported. MO*Ov*POP*surveillance* also has a monthly time step and is simulated for one time step only.

#### 3.3 Process overview and scheduling

##### Processes

Processes included in these models are all related to deer: individual growth (aging), male and female yearling dispersal, bachelor group formation, doe social group formation, fawning, and hunting and non-hunting mortality.

MO*Ov*POP*surveillance* implements three processes individual growth (deer age by one month), non-hunting mortality and hunting mortality with CWD testing.

##### Schedule

Individual growth (increase in age by one month) is scheduled at the beginning of each time step and is followed by non-hunting mortality. This is followed by time step specific processes: bachelor group formation (month = 1), male and female yearling dispersal followed by fawning (month = 5), male yearling dispersal (month = 11) and harvest (month = 12). Group size is updated after any group member executes birth, death or dispersal process. Census and plotting graphs (observer actions) are scheduled at the end of each time step. The month counter resets after every 12 months.

MO*Ov*POP*surveillance* first simulates disease coverage in the pre-harvest population snapshot obtained from MO*Ov*POP, then executes two processes: 1) non-harvest mortality and 2) harvest and sampling for CWD.

The order of processes occurring in MO*Ov*POP and MO*Ov*POP*surveillance* is illustrated in Figure 1.

**Fig 1.**
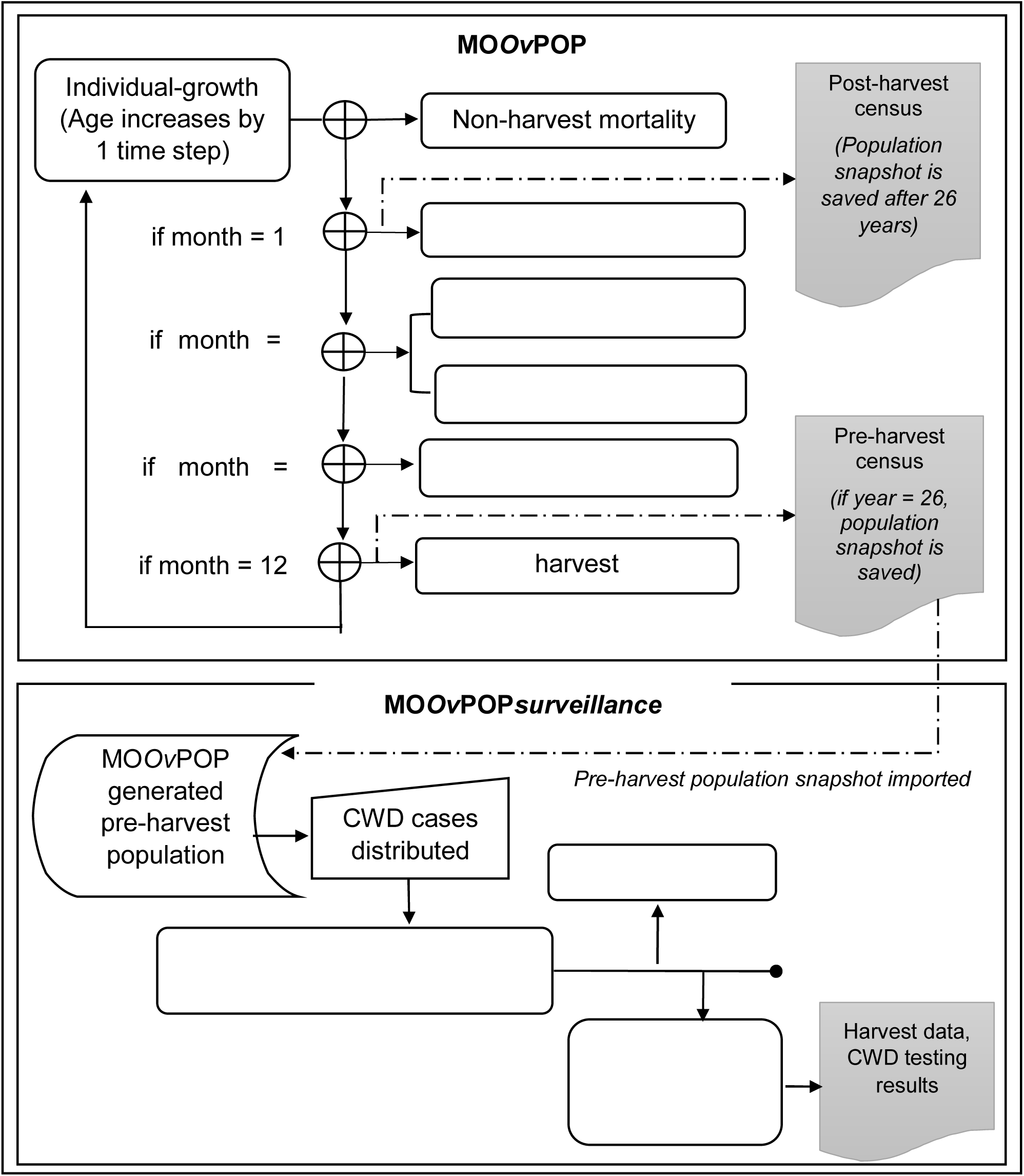
Schedule of processes in MO*Ov*POP and MO*Ov*POP*surveillance*. The rectangular boxes represent actions performed by deer (‘agents’), and the shaded boxes represent actions performed by the model at the population scale.

#### 3.4 Design concepts

##### Emergence

Age and sex structure of the model deer population, as well as the deer distribution pattern, emerge from the model.

##### Adaptation

Fawns (both male and female) entering the yearling class make a decision whether to disperse and leave their natal group. Members of doe social groups update and regulate their group size in response to mortality or birth events involving group members. Group leadership is transferred if a group leader dies.

##### Sensing

Agents (deer) are modeled to ‘sense’ their environment (patch variables or state variables of other agents) before making some behavioral decisions. Yearling bucks perceive the percent forest cover of their home range and determine dispersal distance (described in Submodels: Yearling male dispersal). Doe social group members can sense current group size, group leaders can sense solitary female deer on their patch, and solitary female deer can sense the number of doe social groups and group leaders in their neighborhood (own patch and eight neighboring patches, nine square mile area).

##### Stochasticity

Each individual deer in the model is subject to a mortality probability during a time step. The mortality probability is derived from the mortality rate (natural or hunting). Male and female group leaders are selected from a set of potential candidates during the setup, or when leaders have to be replaced due to mortality events. In MO*Ov*POP*surveillance*, user-specified age-sex prevalence and distribution parameters inform the disease pattern: CWD+ status is assigned to randomly selected deer (random disease distribution) or it is assigned to deer occurring on a randomly selected cluster of patches (clustered disease distribution).

##### Observation

MO*Ov*POP has a graphical display of deer distribution in the landscape. Deer abundance and distribution are updated as the model executes. Additionally, three graphical displays are included: one plots deer abundance versus time, and the other two plot frequency distribution of bachelor and doe social group sizes respectively. Monitors display number of deer in each age-sex-class. Pre-harvest and post-harvest population abundance by age-sex class is recorded in an output file for each year of model run.

MO*Ov*POP*surveillance* interface has monitors for the following outputs: total number of adult deer (male and female) in the population, number of CWD+ deer in the population, number of CWD+ deer in the harvest, total sample size as well as age-sex group wise sample distribution, and CWD+ deer in the sample. For scenarios with clustering of CWD, the graphical display highlights area where CWD cases are clustered in the model landscape. Additionally, following information is recorded in an output file for each year of the model run: number of fawns, yearlings and adults harvested (separately for both sexes), number of CWD+ deer in the hunter harvest, number of CWD+ deer detected in the sample, CWD detection probability and observed (sample) prevalence.

#### 3.5 Initialization

MO*Ov*POP is initialized with GIS coverage data (forest cover) and post-harvest deer density for the desired sampling region. Forest cover data (United States Geological Survey 1992 National Land Cover Data) for selected sampling regions in Missouri was converted to a forest percentage grid of 1 square mile patches to facilitate import in NetLogo. Model deer population for a sampling region is initialized using post-harvest density (deer per square mile), age composition (fawns: yearlings: adults), M:F ratio and age-sex mortality parameters specified by the user. MO*Ov*POP*surveillance* is initialized by importing model-generated pre-harvest deer population snapshot (abundance, sex-age composition and distribution in the landscape) for the sampling region. MOOvPOP and MOOvPOPsurveillance both have NetLogo interface widgets (sliders, choosers, switches) that can be used to set / select user-specified parameters (Figure 2).

**Fig 2.**
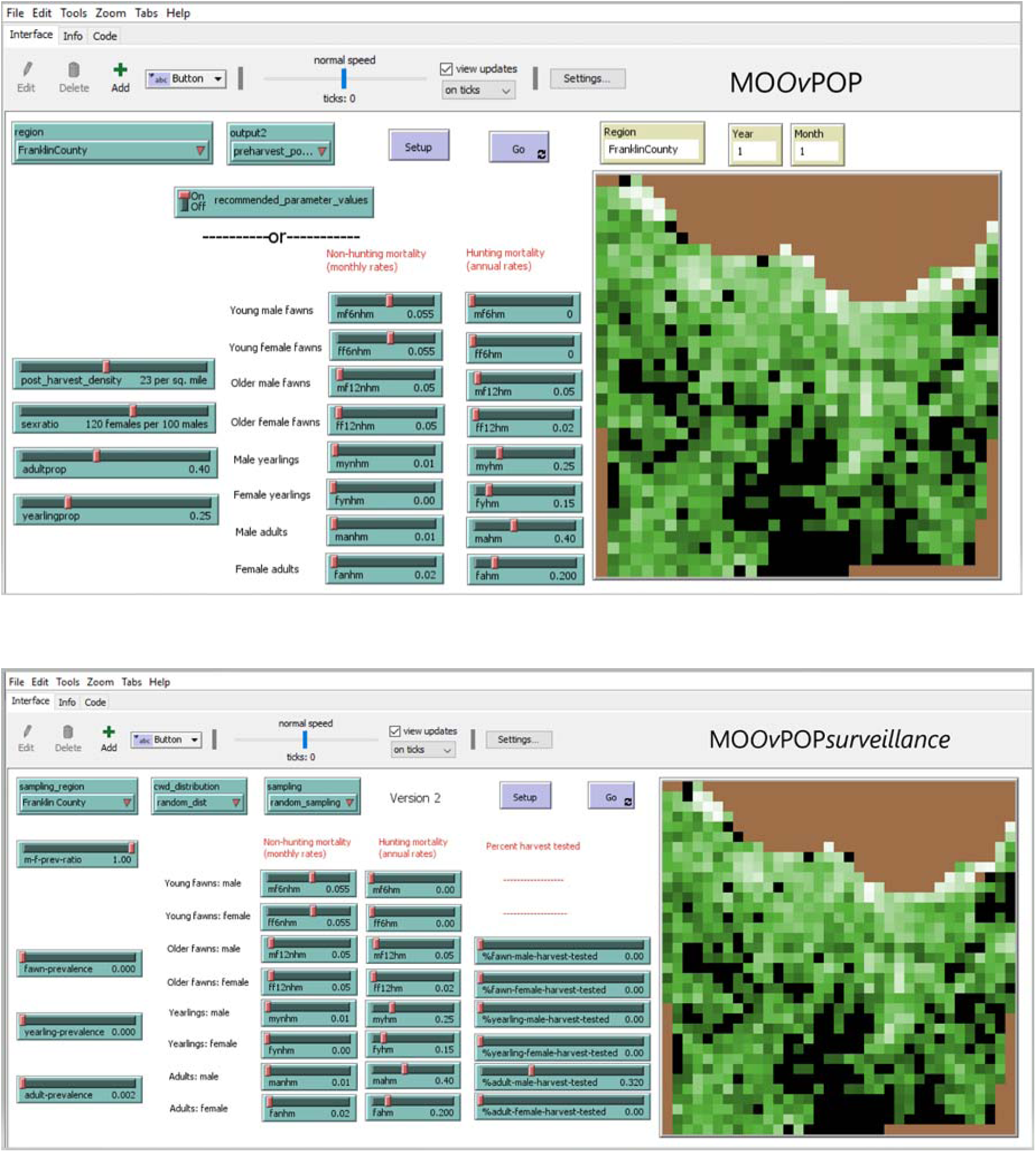
MO*Ov*POP and MO*Ov*POP*surveillance* model interface setup for Franklin County Missouri. Three types of NetLogo interface widgets are provided to set user-specified parameters: sliders, choosers and a switch.

### 3.6 Input data

The model does not use input data to represent time-varying processes.

### 3.7 Submodels

#### 3.7.1 Individual growth

This submodel is executed at the beginning of every time step. All deer in the model landscape update their state variable *aim* (age in months) by one month.

#### 3.7.2 Deer census

Post-harvest census is scheduled in the 1^st^ month (one time step after annual harvest), and pre-harvest census in the 11^th^ month (one time step before the annual harvest) of each year. Number of male and female deer in each age-class is reported separately.

#### 3.7.3 Bachelor-group-formation

Adult male deer are solitary during breeding season, but otherwise form temporary bachelor groups of nonrelated individuals [27]. During the first month every year immediately after post-harvest census, potential number of bachelor groups in the deer population is calculated based on total number of adult and yearling males in the population and the parameter *mean-bachelor-group-size*.

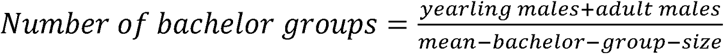

Surviving bachelor group leaders from the previous year maintain their leader status. If the potential number of bachelor groups exceed the available number of group leaders, an appropriate number of bucks older than 32 months are randomly selected and designated as potential bachelor group leaders. The leaders then form bachelor groups by first setting their potential group size (using the parameter *mean-bachelor-group-size*), and then recruiting available adult bucks from patches within a 1.5 mile radius; surviving group members from the previous year are recruited before new members.

#### 3.7.4 Yearling dispersal

The proportion of dispersing yearlings is set using two parameters: *yearling-male-dispersal-rate* and *yearling-female-dispersal-rate*. Dispersing individuals travel the calculated dispersal distance (described below) as an equivalent number of patches in a random direction. We assume that the number of individuals dispersing out of the model landscape is equal to the number of individuals dispersing into the model landscape. Therefore, at any point during dispersal, if a deer moves past the edge of the model landscape (world wraps horizontally as well as vertically), it reappears on the opposite edge as a different deer (its state variable *momid* is changed to 0).

##### Yearling male dispersal

Dispersal distances for yearling bucks are modeled using percent forest cover, as suggested by Diefenbach et al. [39]. Average forest cover for a patch and its immediate neighbors is first calculated for each non-border patch and the value is stored as a patch variable (*dfp*). Border patches set their *dfp* equal to that of one of their non-border neighboring patches. Mean dispersal distance is predicted using the equation from [40]

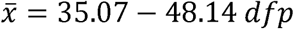

where *dfp* is the patch variable representing mean percent forest of the patch and its neighbors. Variance of dispersal distance is predicted using the equation from [39]

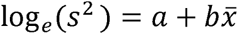

where *a* = 3.51 (SE = 0.597) and *b* = 0.77 (SE = 0.025). Dispersal distance is obtained from a log-normal distribution using the predicted mean dispersal distance and predicted variance of dispersal distance [39]. The dispersal distance is converted from kilometers to miles. If a male yearling reaches a non-deer occupancy patch after dispersal, it is transferred to the nearest deer occupancy patch. After dispersal, the dispersing individual’s state variable *mgroid* takes a value of −1. Group size of the dispersing deer’s natal group is updated.

##### Yearling female dispersal

Dispersal distance for dispersing yearling female is derived from a random distribution using parameters *mean-female-dispersal-distance* and *stddev-dispersal-distance*. If a dispersing individual reaches a non-deer occupancy patch after dispersal, it is transferred to the nearest deer occupancy patch. Dispersing yearling females change their state variables *groid* and *gr* to −1 and −2 respectively. Group size of the dispersing deer’s natal group is updated.

#### 3.7.5 Fawning

A proportion of yearling does aged 13 months (determined by the parameter *juvenile-pregnancy-rate*), and a proportion of adult does (determined by the parameter *adult-pregnancy-rate*) are randomly selected to produce fawns (using ‘hatch-deer’ to create new deer). Yearling does give birth to one fawn and adult deer give birth to twins (MDC data). Sex ratio at birth is set at 1:1 [41]. Fawns inherit two state variables from their dam: *groid* (female social group identifier) and *gr* (group size). Additionally, male fawns have the state variable *mgroid* set to −2.

After the fawning season (month = 5), doe social groups update, and if necessary, regulate their group size. If the group size is greater than 6 (value set by the reporter *doe-social-group-size-regulator*; see Parameterization and Calibration), up to two female group members (adults or yearling) along with their fawns lose group affiliation and become solitary. A deer is considered a member of a doe social group when its state variable *groid* has the group leader’s ID number, and the other state variable *gr* has a value of −1. Designated leaders of doe social groups with four or less members increase their group size by seeking solitary females in a 1.5 mile radius and adding up to two females along with their new-born fawns to the group.

#### 3.7.6 Deer mortality

If a female group leader dies (hunting or natural mortality), leadership is a) transferred to another adult female in the same group (new leader’s state variable *gl* changes from 0 to 1; members change their state variable *groid* to the new leader’s ID (‘who number’); b) if no adult female member exists in the group, surviving members join another group on the same patch with group size <= 3 (change their state variable *groid* to the new group leader’s ID); c) if no small group is available on their patch, the surviving group members become solitary (change their state variables *gr* and *groid* to −2 and −1, respectively).

If members of a doe social group die during a time step, the group leader’s state variable *gr* (accounting for the group size) is decreased accordingly. If there are no members remaining in a group, the leader becomes a solitary deer (state variables *fgroid* and *gr* changed to −1 and −2 respectively).

If a bachelor group leader dies due to natural mortality, leadership is transferred to one of the surviving group members. If there are no members remaining in a bachelor group, the leader changes its status to solitary (state variable ml changed to 0). Sliders are provided on the interface to set age-sex specific mortality rates (Figure 2).

##### Non-hunting mortality

The probability of a deer dying of natural or other non-hunting related causes during every time step is determined by age- and sex- specific monthly mortality rates [42, 43]. Irrespective of these rates, old deer (>240 months) have an overall high probability of dying (0.8) during a time step. Fawns are functional ruminants at two months of age [44], and therefore can possibly survive the death of their mother. We assume that fawns less than two months old do not survive if their mother dies.

##### Hunting mortality

Hunting is the leading cause of deer mortality in most areas of the Midwest [45]. The largest portion of the annual harvest happens during the firearms portion of the deer harvest (usually scheduled between mid- to late November until the 1^st^ week of January), and accounts for most of the samples collected for CWD testing. In this model, harvest is simulated to occur in the 12^th^ time-step every year, one time step after the rut period. Number of deer harvested is specified by age- and sex-specific hunting mortality rates derived from hunter-harvest data collected by the MDC. In MO*Ov*POP, deer surviving the monthly non-hunting mortality are randomly selected to execute the hunting mortality submodel. In MO*Ov*POP*surveillance*, harvest and sampling for CWD can be simulated as a random or non-random process. Sample size for each age-sex class are set using sliders on the interface (*%fawn-male-harvest-tested*, *%fawn-female-harvest-tested*, *%yearling-male-harvest-tested*, *%yearling-female-harvest-tested*, *%adult-male-harvest-tested* and *%adult-female-harvest-tested*). If random harvest and sampling is selected, deer surviving the monthly non-hunting mortality have equal probability of being hunted, and harvested deer are sampled randomly for CWD testing. The assumptions underpinning non-random harvest and sampling processes are derived from the hunter and landowner survey data collected by MDC. The non-random harvest scenario assigns differential harvest probabilities to patches in the model landscape. Twenty five percent of the deer occupancy patches are designated as no-harvest patches, 15 % as high harvest patches and the remaining as low harvest patches. First, no-harvest patches are randomly selected, then from the remaining deer occupancy patches, patches with most deer (average deer density) are selected as high harvest patches. The remaining deer occupancy patches are designated as low harvest patches. Fifty percent of the total annual harvest is set to occur on high harvest patches. Non-random sampling of harvested deer is simulated by assigning differential probabilities for sampling. For every 8000 deer in the population, one high harvest patch is designated as a location for a sampling station. Proximity to a sampling station increases the probability of a harvested deer to be included in the subset that is tested for CWD.

The following counters for each age class-sex category are updated after the harvest: total deer harvested, total CWD+ deer harvested, total deer tested, total CWD+ deer tested.

#### 3.7.7 Disease distribution

Three submodels (*distribute-CWD-fawns*, *distribute-CWD-yearlings*, *distribute-CWD-adults*) select and designate deer as CWD+ during the initial setup of MO*Ov*POP*surveillance*. The selection of deer is informed by user-specified prevalence and disease distribution parameters. Two types of NetLogo interface widgets are provided to set these parameters: sliders (*fawn-prevalence*, *yearling-prevalence*, *adult-prevalence*, and *m-f-prev-ratio*) and a chooser (*cwd_distribution*’). The sliders allow the user to set age-sex class wise prevalence, while the chooser provides the user two choices as a drop down menu – random distribution or clustered distribution of CWD+ deer in the model landscape. Infected individuals (number derived from the prevalence set by the user) are distributed randomly in the model landscape (random distribution) or are clustered in space around a randomly selected deer habitat patch (clustered distribution). For prevalence values < 1%, the area of cluster is ∼5% of the deer habitat. In this model, clustering of CWD cases is best simulated for very low prevalence scenarios when infected individuals are clustered near the point of introduction [18–20].

### 3.8 Parameterization and simulation experiments

#### 3.8.1 Parameterization and Calibration

User-specified information (landscape, vital rates, harvest rates, disease prevalence and distribution) underpins model simulations and dynamics in both the models. We initialized MO*Ov*POP with GIS coverage data (percent forest cover) for Franklin County in eastern Missouri. The model landscape is 35 x 35 patches (∼1225 square miles), of which 951 patches represent Franklin County. Franklin County is of interest because of recent detections of CWD-positive deer, and had the largest opening weekend deer harvest (n=2041) in Missouri during the 2016-2017 deer harvest season [17]. A switch (NetLogo interface widget that can be switched on or off) named ‘*recommended_parameter_values*’ can be toggled to ‘On’ to reset all parameter sliders to values calibrated for sampling regions in Missouri.

Population parameters used to initialize the model deer population, simulate deer behavior or validate model outputs were derived from peer-reviewed literature, field-based surveys, harvest data collected by MDC, or are based on expert opinions (Tables 2 and 3). During initial setup, the abundance and the structure of deer population is determined by four user-specified population parameters: *post_harvest_density*, *sexratio*, *adultprop* and *yearlingprop*. The proportion of fawns in the initial population is calculated by subtracting the sum of *adultprop* and *yearlingprop* from 1 (proportion of fawns = 1 – [*adultprop* + *yearlingprop*]). Here, we initialized MO*Ov*POP with a post-harvest deer density of 23 deer per square mile, a M:F ratio of 1: 1.2, and age composition set at 35% fawns, 25% yearlings and 40% adults.

**Table 2.** Age- and sex-specific mortality parameter values used in MO*Ov*POP and MO*Ov*POP*surveillance*. These values are derived from harvest data collected by the Missouri Department of Conservation.

| Age-Sex Class | Non-hunting mortality <sup>a</sup> |  | Hunting mortality <sup>b</sup> |  |
| --- | --- | --- | --- | --- |
|  | Parameter | Value | Parameter | Value |
| male young fawns (0 – 6 months) | *mf6nhm* | 0.055 | *mf6hm* | 0 |
| female young fawns (0 – 6 months) | *ff6nhm* | 0.055 | *ff6hm* | 0 |
| male older fawns (7 – 12 months) | *mf12nhm* | 0.05 | *mf12hm* | 0.05 |
| female older fawns (7 – 12 months) | *ff12nhm* | 0.05 | *ff12hm* | 0.02 |
| male yearlings (13 – 24 months) | *mynhm* | 0.01 | *myhm* | 0.25 |
| female yearlings (13 – 24 months) | *fynhm* | 0.00 | *fyhm* | 0.15 |
| male adults (> 25 months) | *manhm* | 0.01 | *mahm* | 0.40 |
| female adults (> 25 months) | *fanhm* | 0.02 | *fahm* | 0.20 |

<sup>a</sup> per month; <sup>b</sup> per year

**Table 3.** Population parameters derived for Franklin County deer population from peer-reviewed literature, field-based surveys or expert opinions.

| Parameter | Description | Value | Reference |
| --- | --- | --- | --- |
| Population setup and distribution |  |  |  |
| *post_harvest_density* | Initial deer density (per forested sq. mile) | 23 | MDC |
| *sexratio* | Male: female ratio in the population | 1:1.2 | MDC |
| *adultprop* | Adult proportion ($\geq 25$ months) in the population | 0.4 | MDC |
| *yearlingprop* | Yearling proportion in the population | 0.25 | MDC |
| *min-forestcover-percent* | Minimum percent forest cover of a patch for deer occupancy | 0.25 | [21,22] |
| *max-forestcover-percent* | Maximum percent forest cover of a patch for deer occupancy | 0.75 | [21,22] |
| Behavior |  |  |  |
| *mean-bachelor-group-size* | mean (± sd) number of adult bucks in a bachelor group | 2 - 5 | [44,48,49], L.Hansen, pers.obs |
| *doe-group-size-regulator* | doe social group size above which it undergoes fission | 6 | [46,47], L.Hansen, pers.obs |
| *juvenile-pregnancy-rate* | proportion of female fawns that reproduce | 0.2 | J. McRoberts, pers.comm. |
| *adult-pregnancy-rate* | proportion of adult females that reproduce | 0.8 | MDC |
| *yearling-male-dispersal-rate* | proportion of dispersing yearling males | 0.46 | [39,40] |
| *yearling-female-dispersal-rate* | proportion of dispersing yearling females | 0.22 | [51] |
| *mean-female-dispersal-distance* | mean dispersal distance for yearling females (miles) | 11 | [31] |
| *stddev-dispersal-distance* | standard deviation for the mean dispersal distance of yearling females (miles) | 4 | [31] |

Population dynamics of the model deer population is defined by two sets of user-specified age-sex-specific parameters, *hunting mortality rates* and *non-hunting mortality rates*. Hunting mortality rates are annual, while non-hunting mortality rates are monthly, and the parameters for Franklin County used here were derived from harvest data collected by MDC (Table 2).

Group sizes in the model are regulated using reporters, *mean-bachelor-group-size* and *doe-group-size-regulator*. In NetLogo, a reporter is a set of instructions for computing a value. Parameters used in *mean-bachelor-group-size* and *doe-group-size-regulator* were calibrated so that the model group sizes remained within the range derived from expert opinion (L. Hansen, pers obs) and peer-reviewed literature [44,46–49].

Body mass attained during the breeding season appears to be a strong determinant of a fawn’s ability to breed [50]. The percent of female fawns that breed is influenced by the population’s level of nutrition. The reporter, *juvenile-pregnancy-rate*, was set at 20%, based on data from an ongoing deer study in Missouri (Jon McRoberts, personal communication). The proportion of adult females that breed, *adult-pregnancy-rate*, was set at 80% (derived from MDC data https://extension2.missouri.edu/g9479).

Dispersal rates for yearling bucks range between 46 and 80% [40] but predictive relationships are poorly understood [39]. We have set the yearling buck dispersal rate at 46% for simulations described in this paper. Similarly, based on observations from west-central Illinois, the yearling doe dispersal rate is set at 22% [51]. Mean dispersal distance for yearling does was set at 11 miles – this value was extrapolated for a post-harvest deer density of ∼25 per forested km^2^ from a logistic regression model based on meta-analysis of yearling doe dispersal data [31].

## 4. Model evaluation

### 4.1 Evaluation of MO*Ov*POP performance

First, we initialized MOOvPOP with vital rates derived for Franklin County deer population and simulated population growth over a period of 25 years. We completed five iterations to assess deer population trends. Age-sex class composition was plotted throughout each model run and annual finite population growth rate (λ) was determined using pre-harvest abundance from year two onwards. The age-sex class composition of model-generated deer populations was stable after five years. Results for one of the model iterations are presented in Figure 3. The finite population growth rate equilibrated to 1 after year 15, indicating a stationary model deer population (average λ ∼ 1) (Figure 4). MDC’s county-specific deer population assessment for 2016 supports the stable deer population trend for Franklin County (Missouri Deer Season Summary and Population Status Report for 2017). Therefore, it is advisable to run MOOvPOP for at least 15 years before obtaining a pre-harvest population snapshot. We selected the 26^th^ year pre-harvest population snapshot for initializing MO*Ov*POP*surveillance*.

**Fig 3.**
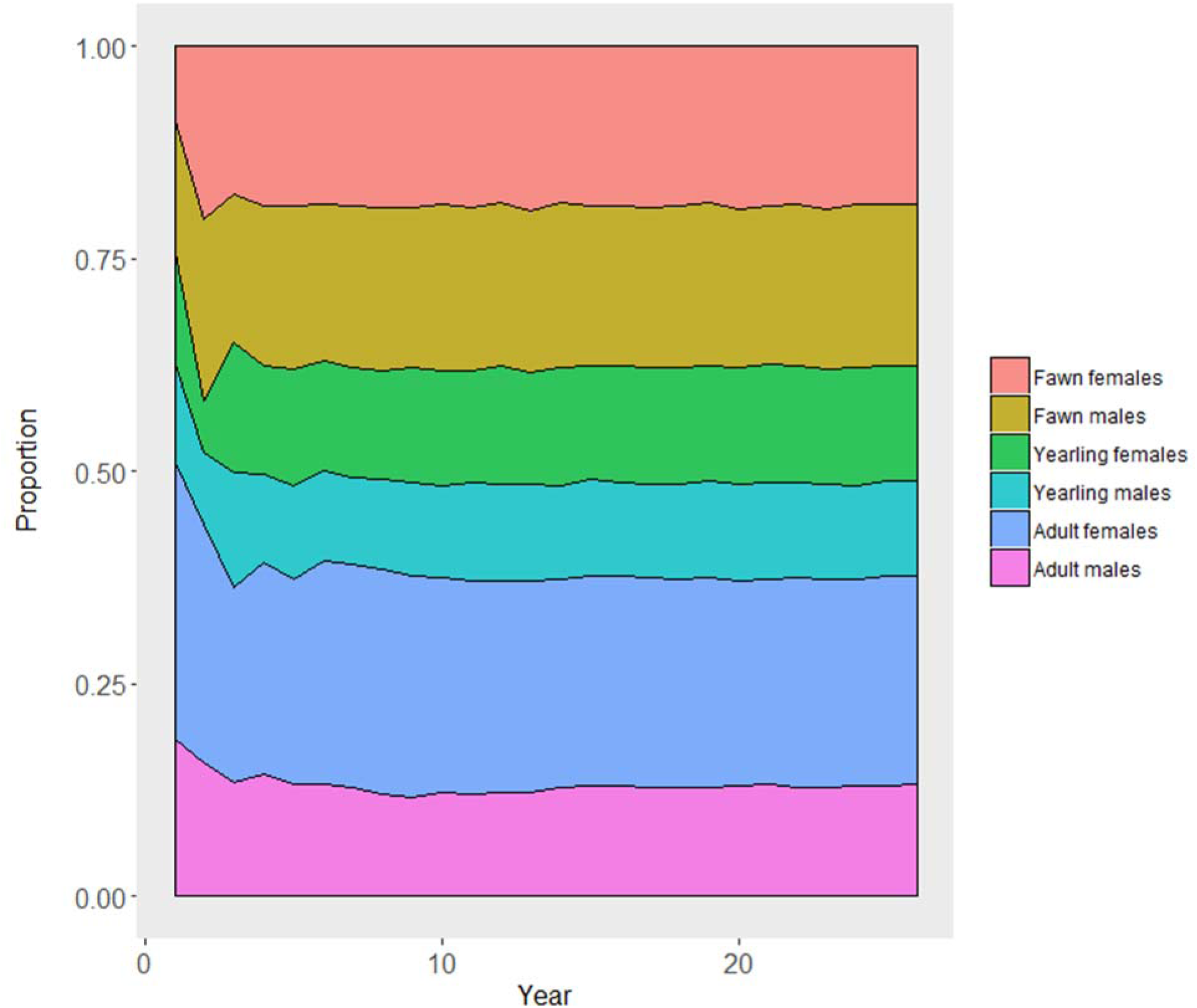
Age-sex composition of MO*Ov*POP simulated Franklin County deer population over a period of 25 years. The age and sex composition of the model population remained stable after the 5^th^ year.

**Fig 4.**
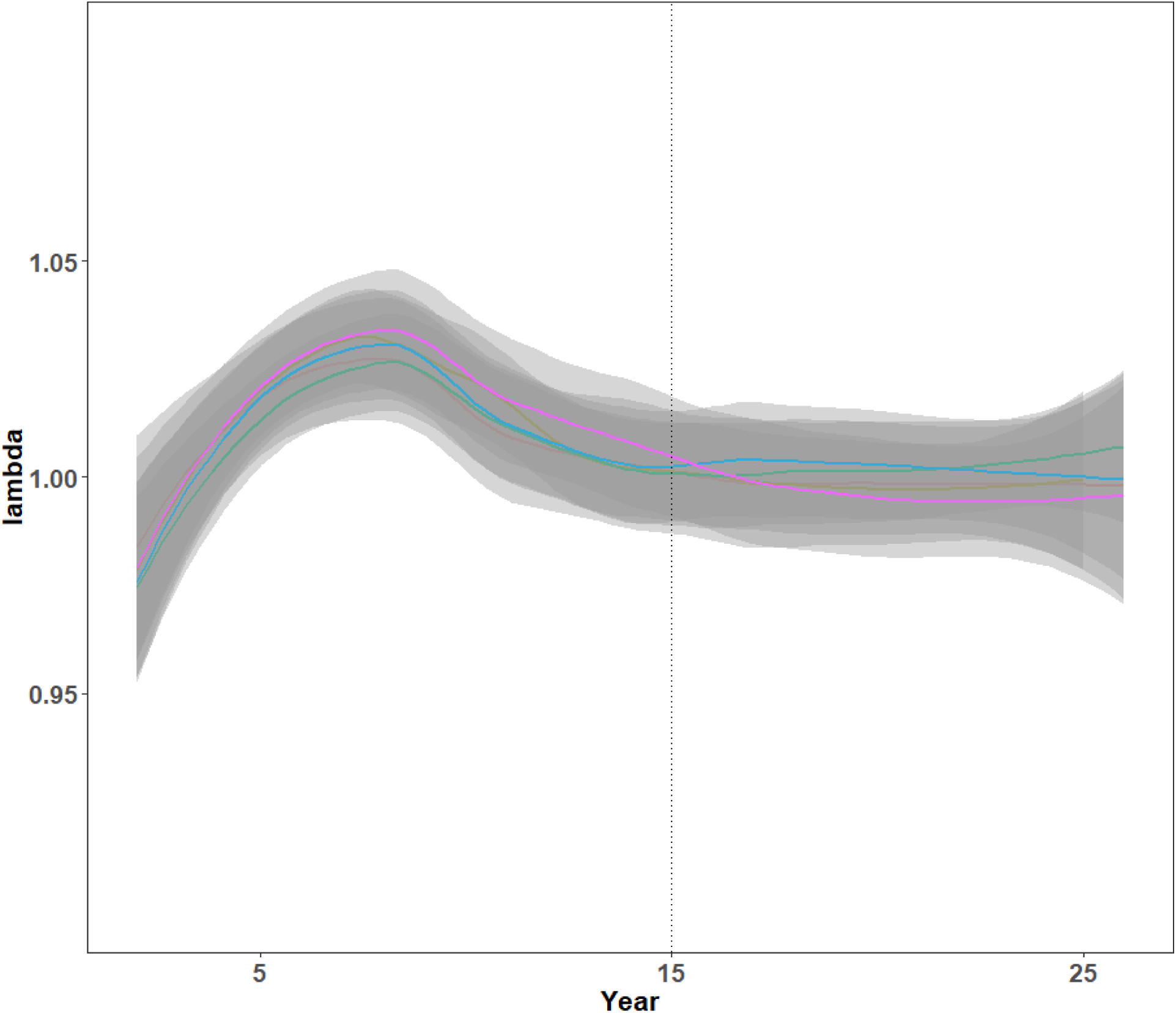
Population growth rate (λ) for the Franklin County, Missouri, deer population simulated using MO*Ov*POP (five iterations). The population reached a relatively stationary abundance after 15 years (vertical line), when λ converged to 1.

We then completed 100 iterations of MOOvPOP and analyzed pre-harvest deer population snapshots obtained in the 26^th^ year of model run. For all model iterations, group sizes remained between two and 11 for doe social groups and between two and eight for bachelor groups. Group size distribution for one of the model-generated deer population snapshot is presented in Figure 5. The group size distribution is in agreement with deer group sizes reported in literature and expert opinions: Doe social group size ranges between two and 12 [46, 47], but smaller group sizes (less than eight) are commonly seen in Missouri, and bachelor group size typically ranges between two and five [44,48,49], but larger groups of up to eight are occasionally seen in Missouri (L. Hansen, personal observation). The mean pre-harvest deer abundance for the 100 population snapshots was 25, 524 (ranging between 23,137 and 28,735), while MDC’s Cervid Program estimated pre-harvest population for Franklin County in 2016 was 26,502 (MDC unpublished data). MDC’s Cervid Program assesses county-specific deer populations using a population simulation model that incorporates harvest data, age-at-harvest data, and estimated survival and reproduction rates.

**Fig 5.**
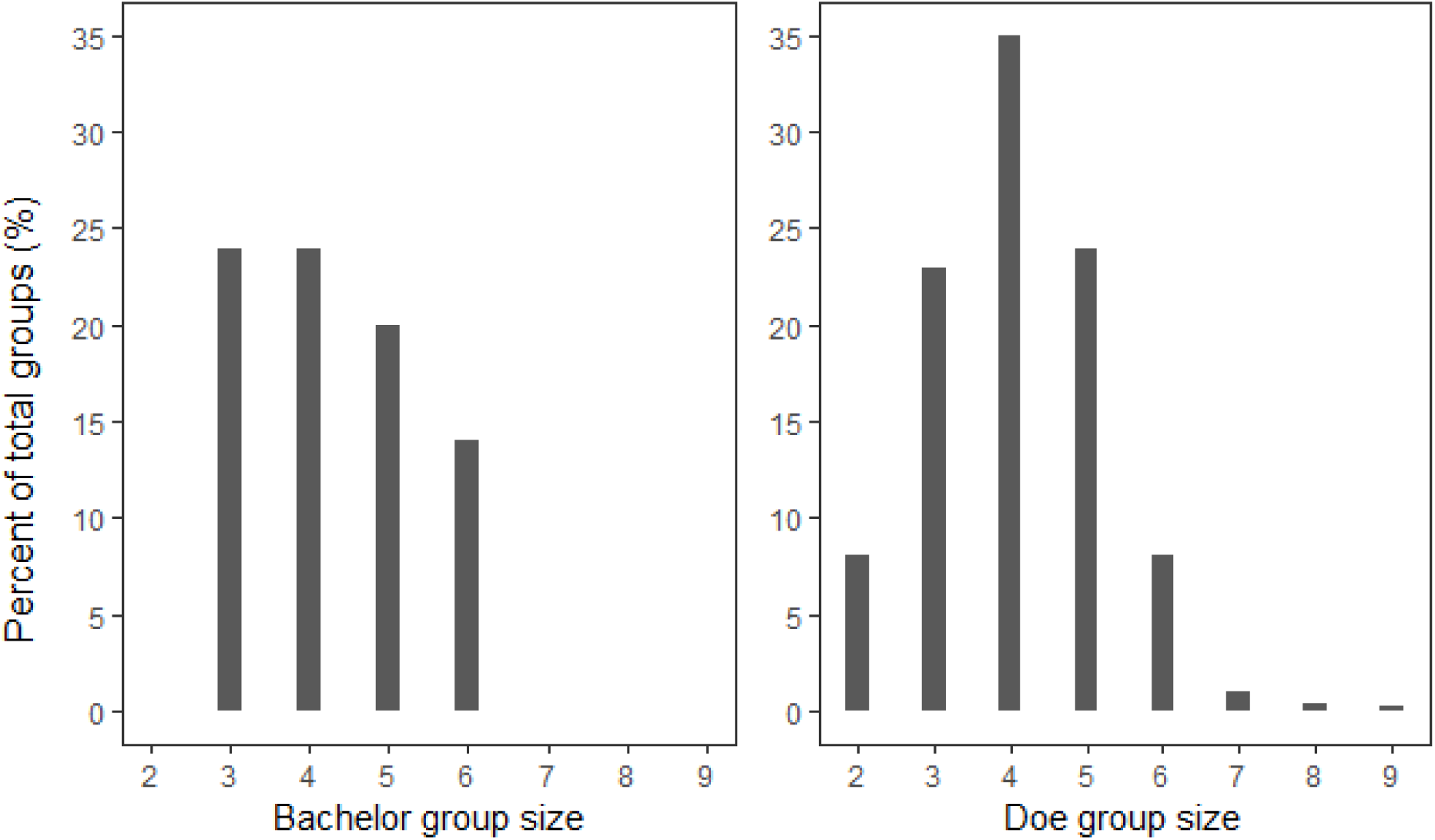
Group size distribution for bachelor and doe social groups in a model-generated deer population snapshot for Franklin County, Missouri. Group size distributions were in agreement with references and expert opinions throughout the model run.

MO*Ov*POP derived deer population snapshots (n=100) were composed of 34% fawns (minimum 33%, maximum 34%), 24% yearlings (minimum 24% maximum 25%) and 42% adults (minimum 41%, maximum 42%). Estimates derived from the 2016-17 deer harvest data for Franklin County are comparable: 35% fawns, 25% yearlings and 40% adults. The M:F ratio for the model-derived deer population snapshots was 1: 1.18 (minimum 1:1.14, maximum 1:1.21), and the estimate derived from Franklin County harvest was 1:1.12.

Our assessments indicate that MO*Ov*POP generated population snapshots are sufficiently realistic for the purpose of simulating harvest and harvest-based surveillance scenarios.

### 4.2 Evaluation of MO*Ov*POP*surveillance* performance

Prevalence-sample size scenarios were assessed using hypergeometric approximation and MO*Ov*POP*surveillance*. Standardized sample size tables and online calculators like EpiTools (Sergeant, ESG, 2018. Epitools epidemiological calculators. Ausvet Pty Ltd. Available at: http://epitools.ausvet.com.au) use the hypergeometric approximation to determine the sample size for disease detection in a population. The underlying assumption for this approximation is that the samples are obtained using a probabilistic method [11]. Canon and Roe (1982) have provided the following equation to determine

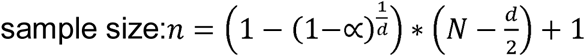

where N is the population size, d is the number of diseased individuals in the population, α is the desired confidence level (or the detection probability) and n is the required sample size. Using this approximation, we calculated sample sizes with high detection probabilities (90%, 95% and 99%) for a population of 4000 deer with CWD prevalence of 0.5%, 1%, 2% and 5%. The same prevalence – sample size combinations were used to simulate MO*Ov*POP*surveillance*. MO*Ov*POP-generated Franklin County deer population snapshot was used to initialize the model, and CWD coverage, harvest and sampling for CWD were simulated as random processes. CWD+ cases were distributed in adult male deer only; sampling for CWD was therefore limited to adult male harvest. The pre-harvest abundance of adult male deer in this model-generated population was 4387. We used this simple scenario to facilitate comparison of detection probabilities between the two methods.

Ten replicates of 100 MO*Ov*POP*surveillance* iterations were undertaken for each prevalence-sample size scenario. For each replicate, detection probability was calculated as the number of iterations in which at least one CWD+ deer was detected. We compared model-derived and hypergeometric approximation derived detection probabilities using a one-sample t-test. Mean detection probabilities derived from model iterations for each prevalence-sample size scenario were no different than the hypergeometric approximation derived detection probabilities (Table 4).

**Table 4.** Comparison of hypergeometric approximation-derived and MO*Ov*POP*surveillance*-derived detection probabilities for hypothesized prevalence-sample size scenarios in a population of 4000 hosts. Disease coverage as well as the sampling process is assumed to be random. Ten replicates of 100 model iterations were used to obtain the mean detection probability for each scenario.

| Prevalence (%) | Detection probability (%)<sup>*</sup> | Sample size <sup>*</sup> | Model-derived mean detection probability (%) (95%CI, t(df), p) # |
| --- | --- | --- | --- |
| 0.5 | 90 | 435 | 89.9 (88.2 – 91.6, t(9)=0.13, p=0.899) |
|  | 95 | 556 | 93.7 (91.6 – 95.8, t(9)=1.41, p=0.191) |
|  | 99 | 822 | 98.8 (98.1 – 99.5, t(9)=0.61, p=0.555) |
| 1 | 90 | 224 | 89.3 (87.8 – 90.8, t(9)=1.07, p=0.31) |
|  | 95 | 288 | 93.8 (91.8 – 95.8, t(9)=1.33, p=0.217) |
|  | 99 | 434 | 99.1 (98.5 – 99.7, t(9)=0.36, p=0.726) |
| 2 | 90 | 113 | 90 (87.7 -92.3, t(9)=0, p = 1) |
|  | 95 | 147 | 94.4 (93.2 – 95.6, t(9)=1.15, p=0.279) |
|  | 99 | 223 | 98.5 (97.5 -99.5, t(9)=1.17, p=0.272) |
| 5 | 90 | 46 | 88.7 (86.9 – 90.5, t(9)=1.62, p=0.140) |
|  | 95 | 59 | 94.6 (93.3 – 95.9, t(9)=0.68, p=0.508) |
|  | 99 | 90 | 99.2 (98.6 – 99.8, t(9)=0.80, p=0.443) |

□ Approximation of the hypergeometric distribution.

### 4.3 Sensitivity analysis

Disease coverage in the population and the manner in which samples are obtained are two critical processes that can influence the outcome of a disease surveillance strategy. It is important to assess how sensitive model outputs are to changes in the way these processes are modeled as our understanding of ‘real world’ patterns of such processes is fraught with uncertainties. We evaluated how MO*Ov*POP*surveillance*-derived detection probabilities responded to changes in a) CWD clustering pattern and b) non-random sampling process.

MO*Ov*POP*surveillance* was initialized with MO*Ov*POP-generated Franklin County deer population snapshot and CWD prevalence was set at 1% in adult deer. The sample size for CWD surveillance was set at 50% of the adult harvest (∼ 1460 harvested adult deer tested for CWD). For each scenario described below, we undertook ten replicates of 100 model iterations.

The baseline scenario did not have any disease clustering (randomly distributed CWD cases), and harvest as well as sampling of harvested deer were simulated as random processes. For the ‘Alternate’ scenario, 100% of the CWD cases were clustered in space, on an area equivalent to ∼5% of the deer habitat. Clustering of cases occurred on a randomly selected group of contiguous patches. Harvest was simulated as a non-random process by designating 15% of deer habitat patches as high harvest pressure patches. Half of the total deer harvest (50%) occurred on high harvest pressure patches. Sampling of harvested deer was also simulated as a non-random process and is influenced by proximity to a sampling station. Figures 6 and 7 illustrate CWD distribution (random and clustered) and sampling (random and non-random), respectively.

**Fig 6.**
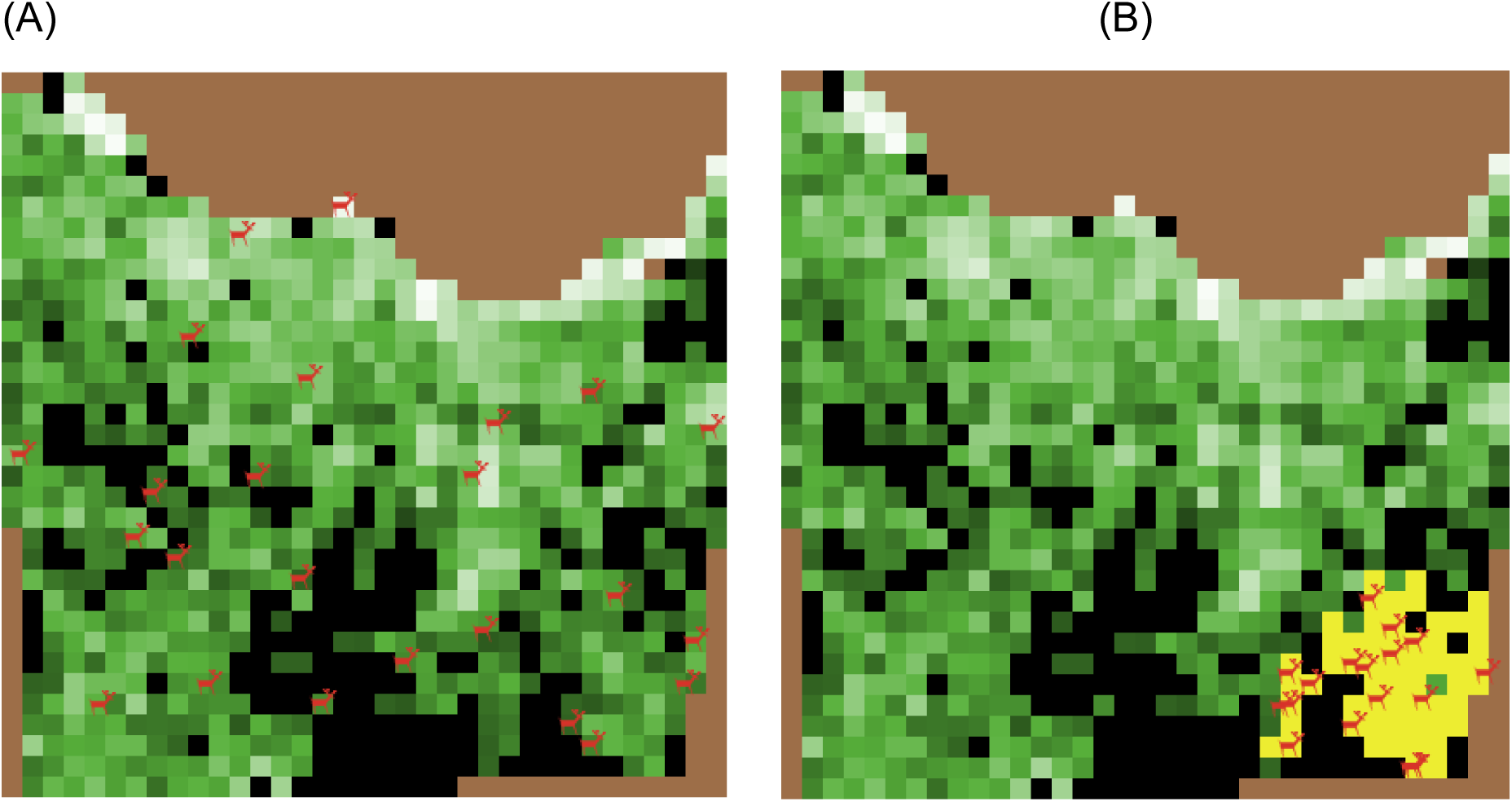
Random (A) or clustered (B) distribution of infected deer in MO*Ov*POP*surveillance* landscape representing Franklin County, Missouri. Infected deer (red) are shown on the model landscape, and yellow colored patches in (B) represent the disease cluster.

**Fig 7.**
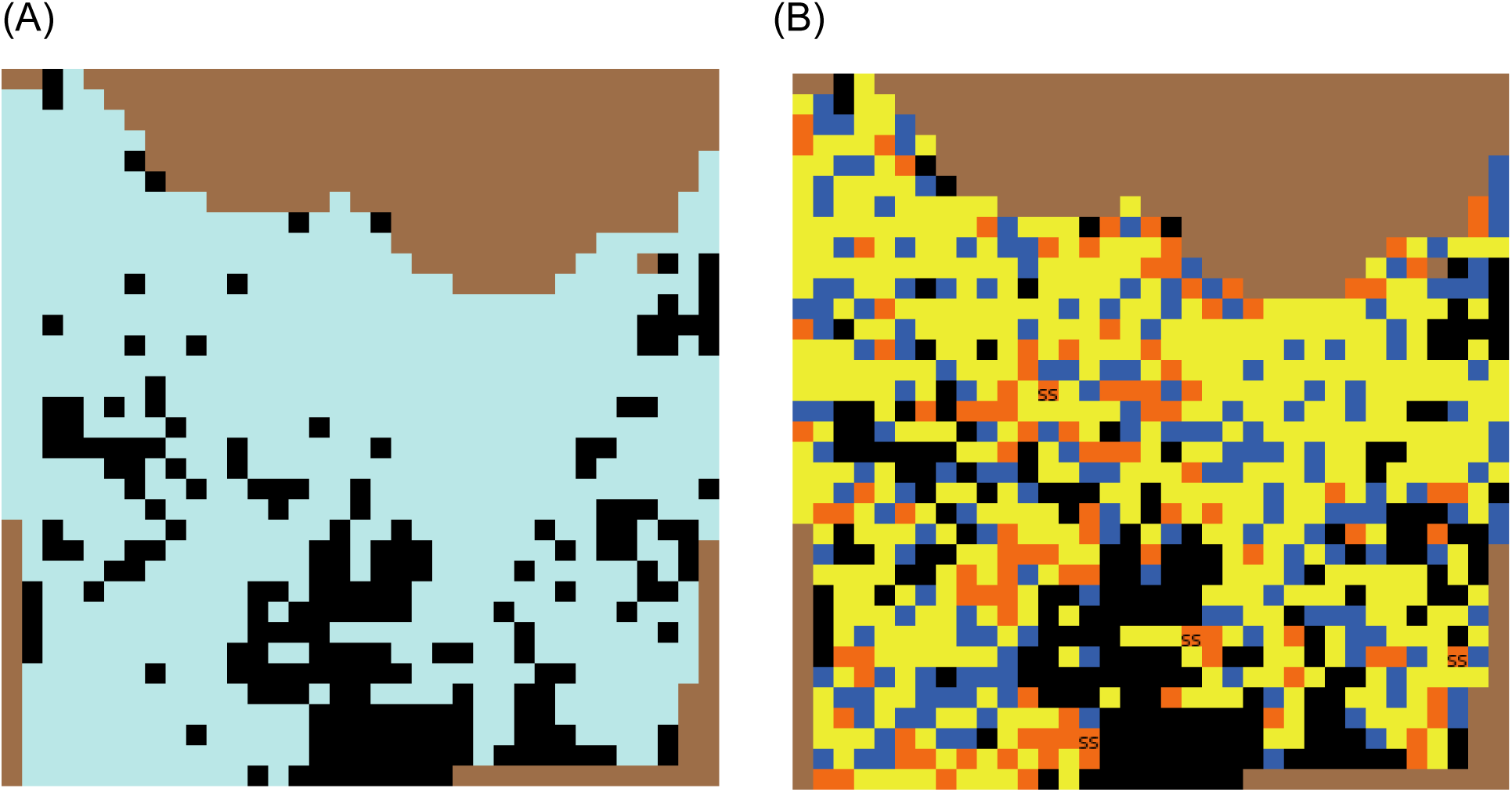
(A) MO*Ov*POP*surveillance* landscape representing Franklin County with uniform harvest pressure. (B) The same landscape with non-uniform harvest pressure to simulate non-random sampling. Blue color indicates patches where deer are not harvested, yellow indicates low harvest pressure and orange indicates high harvest pressure. Patches labelled ‘SS’ indicate sampling stations.

We then explored the effect of changing the CWD clustering pattern. Instead of 100% clustering of cases (Alternate scenario), we simulated patterns with an incremental proportion of CWD+ deer (2%, 4%, 6%, 8% and 10%) randomly distributed outside the cluster (cluster98%, cluster96%, cluster94%, cluster92% and cluster90%). For these scenarios, harvest and sampling were simulated as non-random processes similar to the alternate scenario.

Finally, we explored non-random harvest patterns for their effect on CWD detection probabilities. The proportion of high-harvest patches (15% in the Alternate scenario) was increased incrementally by 3%: ‘non-random sampling 18’, ‘non-random sampling 21’, ‘non-random sampling 24’, ‘non-random sampling 27’, ‘non-random sampling 30’. In these scenarios, percent of harvest occurring on high harvest patches (50%) as well as clustering of CWD cases was similar to the Alternate scenario.

Results of the sensitivity analysis are presented in Figure 8. Without any clustering of CWD cases and random sampling, the baseline scenario consistently yielded high detection probabilities (100%). Detection probability decreased to ∼76% with the alternate scenario, where 100% of CWD cases were clustered and 15% of deer habitat patches had high harvest pressure (non-random sampling). Detection probability was sensitive to changes in the number of infected individuals outside the cluster. For the Franklin County deer population with 1% CWD prevalence in adult deer and a sample size of ∼1460, the confidence in detecting CWD increased from 76% to 90% when 8-10 infected deer occurred outside the cluster. This is particularly relevant for very low prevalence situations, for instance, where CWD is recently introduced and all the cases are clustered around the point of infection. The sample size threshold of detecting CWD confidently would increase later in the course of the epizootic after CWD cases spread to areas outside the cluster. Changing the non-random harvest pattern did not have much effect on the detection probability.

**Fig 8.**
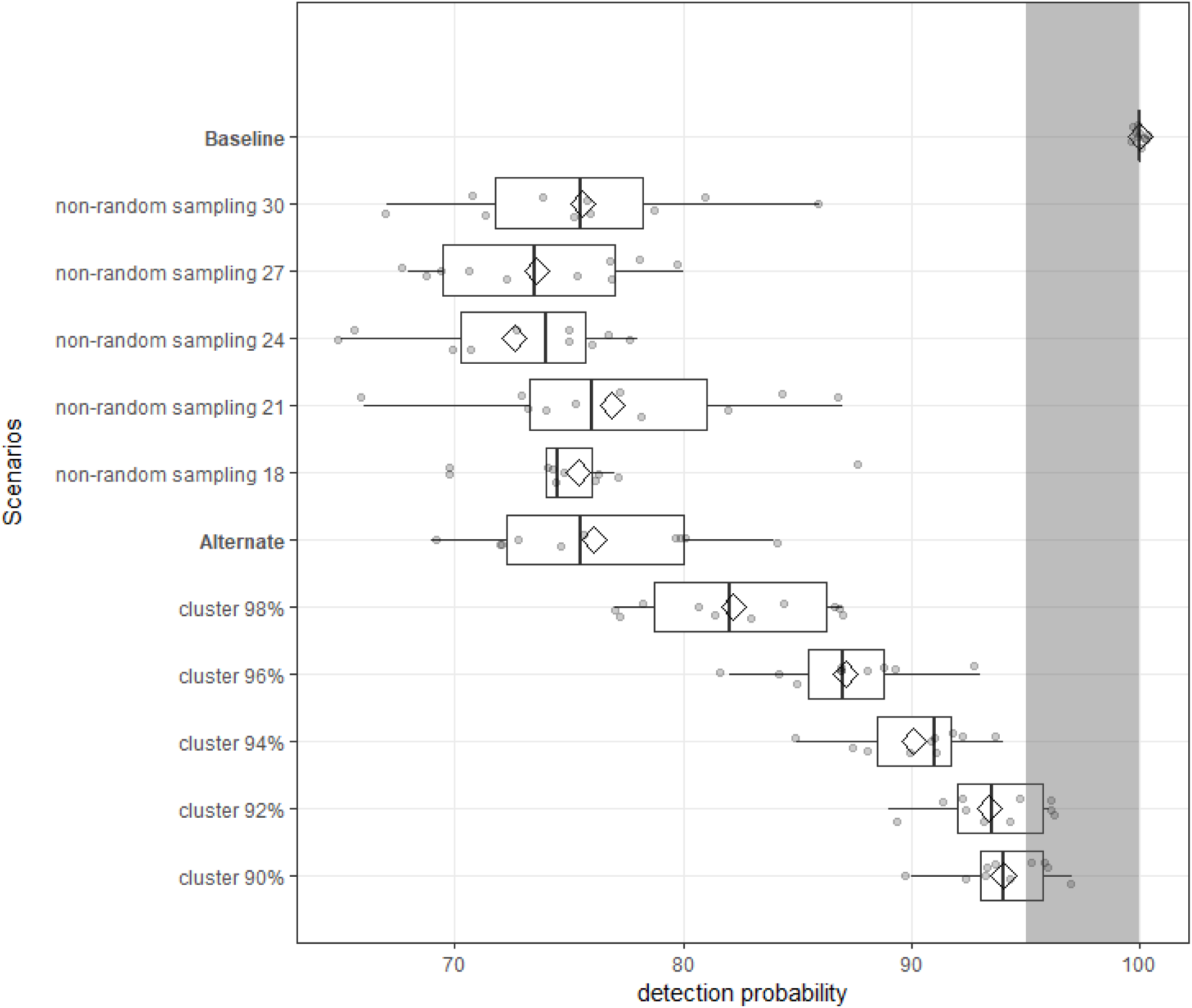
Model-derived detection probabilities for Franklin County deer population with different CWD clustering and/or harvesting (sampling) patterns. CWD prevalence was simulated at 1% (adult deer only) and the sample size was set at ∼1460 (= 50% adult harvest). Non-random sampling scenarios indicate the proportion of high harvest pressure patches in the model landscape and cluster scenarios indicate the proportion of CWD cases occurring within the cluster. Baseline scenario had no clustering of CWD cases and uniform harvest pressure, while alternate scenario had 100% of CWD cases within the cluster and 15% of the patches designated as high harvest patches. Shaded area indicates high detection probabilities (≥ 95%).

## 5. Model Application

We now illustrate how the models can be used to support CWD surveillance. Simulations were designed to estimate the confidence in detecting CWD (detection probability) for a range of prevalence - sample size combinations. We used five hypothesized prevalence levels in adult deer of Franklin County (0.2%, 0.4%, 0.6%, 0.8% and 1%). Sample sizes were simulated as percent of the total adult deer harvested (10%, 20%, 30%, 40% and 50%). The adult deer harvest in our model (∼3000) approximates the adult harvest during 2016 Fall Firearm harvest in Franklin County (MDC Deer Harvest Summary 2016-2017). For each hypothesized prevalence - sample size combination, we simulated baseline and alternate scenarios (described under Sensitivity analysis). We undertook ten replicates of 100 iterations of the model for each prevalence-sample size and scenario combination.

Under the baseline scenario, 17 of 25 prevalence-sample size combinations had high detection probabilities (≥95%) (Figure 9). Though CWD detection probability increased with sample size and CWD prevalence, the relationship is not linear, and even under such ideal assumptions the likelihood of reaching the 95% detection threshold was low for the lowest prevalence rates (typical of the low prevalence rates of CWD that currently are thought to occur in Missouri) and at the lowest sampling efforts. Thus, under ideal assumptions, model-derived sample sizes for confident detection of CWD in Franklin County deer population range between 1189 (CWD detection threshold set at 0.2% prevalence) and 297 (CWD detection threshold set at 1% prevalence). There was a strong increase in sample sizes necessary for >95% detection probability for each scenario as prevalence rates decrease. For example, necessary sampling efforts double as prevalence declines from 1% to 0.6% and increases 3 fold - 5 fold at lower prevalence.

**Fig 9.**
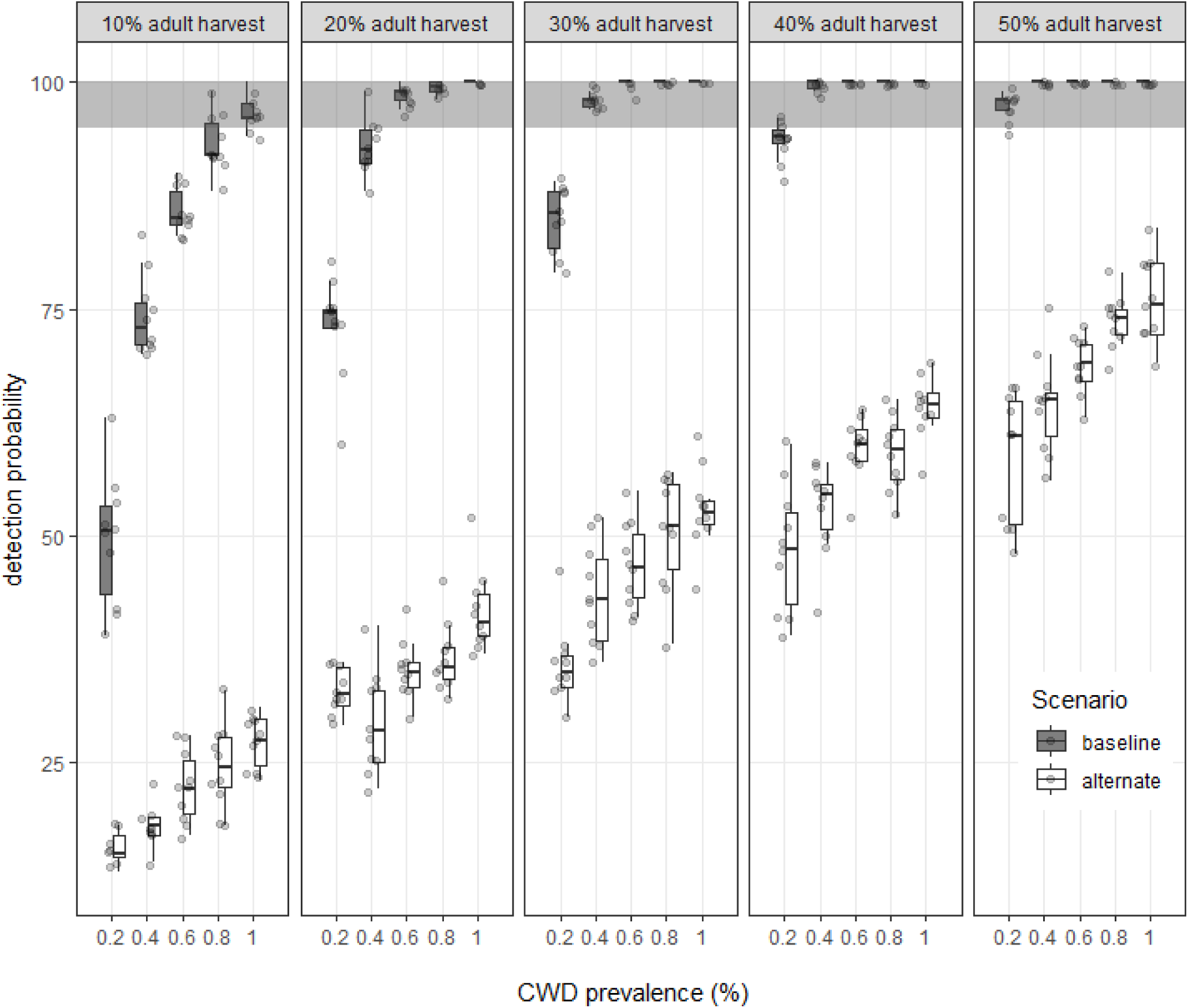
Model-derived detection probabilities for a range of sample sizes (10% to 50% of adult deer harvest) and CWD prevalence levels (0.2% to 1%). The baseline scenario assumes random distribution of infected individuals in the population, random harvest and random sampling. The alternate scenario assumes clustered distribution of infected individuals, non-random harvest and non-random sampling. Shaded area indicates high detection probabilities (≥ 95%).

The alternate scenario with spatial clustering of CWD and biased hunter harvest had strong impacts on CWD detection probabilities. None of the tested sample sizes under the alternate scenario had high detection probabilities. Even with 1% prevalence, the largest sample size that we tested (50% of the adult harvest for Franklin County = ∼ 1500) had a detection probability of 78%. Detection probabilities further decreased as prevalence decreased, and for 0.2% prevalence it dropped to 58%. We explored increasing prevalence levels under the alternate scenario to determine the prevalence threshold at which the sample size ∼ 50% of adult harvest (∼1500 deer tested) has a high detection probability. This sample size approximates total deer sampled from Franklin County during 2016 hunting season and represents the upper limit of sample size under realistic conditions. Under ideal assumptions, a sample size of ∼1500 should be able to confidently detect CWD when the adult prevalence is less than 0.2% (<20 infected adults in the population). Our model explorations indicate that under realistic assumptions, this sample size can confidently detect CWD (at least one CWD+ deer in the sample) when the adult population prevalence is more than 5% (>500 infected adults in the population).

The results of these simulation experiments (scenario-detection probability data) can be used to determine CWD detection thresholds, calculate optimum sample sizes to confidently detect CWD in target populations, and design a sustainable CWD surveillance strategy.

## 6. Discussion

Sampling is a fundamental component of epidemiological surveillance as sample data are used to infer about the presence or absence of a disease in the target population. Surveillance of important wildlife diseases like avian influenza, bovine tuberculosis, brucellosis, West Nile virus, and chronic wasting disease often rely upon samples obtained from hunter-harvested individuals [1,3,4,19,52–56]. But hunter harvest sampling does not approximate random sampling or any other probabilistic sampling method, and such samples are not representative of the population. Nonetheless, wildlife management agencies and researchers often calculate sample size (and associated confidence level) necessary for disease detection in a target population under the assumption that harvest-based sampling approximates probabilistic sampling [14,53,56–58]. Such an approach may result in misleading and biased inferences about the presence or absence of a disease in the target population.

Using computer simulation and simplified assumptions (uniform deer density across a 30 square mile landscape, and CWD cases clustered in the center of the landscape), Samuel et al. [5] demonstrated that non-random sampling decreases the CWD detection probability. Similarly, Nusser et al. [13] used a simulated white-tailed deer population for one county in Iowa, USA to compare probability-based and alternative sampling strategies, and their results similarly demonstrated that sample sizes necessary for confidently detecting CWD are underestimated if assumptions of random CWD distribution and random sampling are used for sample size calculations.

As recommended by Nusser et al. [13], we have developed a model-based tool that can incorporate real world heterogeneities and biases, and can be applied to different regions / deer management units to guide collection and analysis of CWD surveillance data that relies on non-probabilistic methods like harvest-based sampling. An important feature of our agent-based framework is the ability to simulate age-sex specific scenarios and interventions as relevant individual host characteristics (age, sex, group membership) and behaviors (dispersal, grouping behavior) have been incorporated in the model programs. Further, the model programs are designed to be user friendly; the interface sliders and choices allow users to update model assumptions based on their current best knowledge of the system, and thereby employ an adaptive management approach for defensible decision making while planning and implementing CWD surveillance activities.

Results of the simulation experiments described in this paper (Table 5) show that model-derived sample size requirement for detecting CWD with ≥ 95% confidence are lower for the ecologically unrealistic baseline scenario that assumes random sampling and random distribution of infected animals, compared to the alternate scenario that incorporates realistic biases in sampling and disease distribution. For example, if the objective of surveillance in Franklin County is to detect CWD with high confidence (≥ 95%) when 1% (or more) of adult deer are infected, then under the baseline scenario (random CWD distribution and random sampling) the model-derived sample size is ∼ 297 (10% of adult deer harvest for Franklin County). Sample size tables (based on hypergeometric distribution, and assumption of random sampling) also recommend a sample size of ∼300 from a host population (∼adult deer population in Franklin County) with disease prevalence of 1% if the objective is disease detection with high confidence (95%) [5, 11]. But under the alternate scenario with 1% prevalence, the sample size required for detecting CWD with high confidence (≥95%) will be larger than 1500 (∼50% adult deer harvest for Franklin County). Unrealistic assumptions result in underestimation of sample sizes due to the failure to account for spatial structure of the deer population, CWD and harvest.

**Table 5.** Sample sizes with detection probabilities for Franklin County contrasted for the baseline (random CWD distribution, random harvest and sampling) and alternate (clustered CWD distribution, non-random harvest and sampling) scenarios.

| **Scenarios** | **Prevalence** |  |  |  |  |
| --- | --- | --- | --- | --- | --- |
|  | **0.2%** | **0.4%** | **0.6%** | **0.8%** | **1%** |
| Random CWD distribution,<br>random sampling | 1189<br>(≥95%) | 891<br>(≥95%) | 594<br>(≥95%) | 297<br>(≥95%) | 297<br>(≥95%) |
| Clustered CWD distribution,<br>non-random sampling | 1486<br>(59%) | 1463<br>(64%) | 1463<br>(75%) | 1463<br>(75%) | 1463<br>(78%) |

The importance of determining sample size targets using realistic assumptions cannot be overemphasized, especially if the main objective of surveillance is to detect the disease in areas where it has not been found yet so that control measures may be rapidly implemented. As CWD is known to occur at a low prevalence near the point of introduction during the early stages of the epizootic, sample size targets should be determined using a low prevalence detection threshold [18–20]. The trade-off between sample size and CWD detection threshold is an important consideration: the lower the detection threshold is set, the larger the required sample size to confidently detect CWD in the target population. But sampling effort also has to be considered in the context of sustainability of surveillance efforts, as resources to support sampling efforts are limited and subject to waning public interest. On the other hand, the very purpose of surveillance is defeated if the detection threshold is set at a higher prevalence level to facilitate use of a lower sample size. Our modeling framework can be used to optimize surveillance strategies by 1) applying the model to smaller units (for example, township units) to ensure reasonable sample size, low detection threshold, and meaningful inferences about CWD occurrence and distribution; and 2) developing a ‘risk map’ based on sampling history and other factors like proximity to areas where CWD has been detected, captive facilities, geographic features, etc. to prioritize areas for sampling.

Sampling history for a region (previous sampling effort and outcomes) can be useful for making valid inferences about the presence or absence of the disease in the local population. In particular, negative surveillance outcomes (no infected individuals detected in the sample) can be used to infer the upper limit for the number of infected individuals that could be present in the population. But such inferences would be misleading if the sampling method assumes random sampling when surveillance activities are based on non-random sampling. For instance, if no CWD-infected deer are detected in a random sample of ∼1189 adult deer harvested from Franklin County, then the inferred upper limit of CWD prevalence in Franklin County can be estimated at 0.2% (∼ 20 infected adult deer) (Table 5). But under realistic assumptions (clustered CWD distribution and non-random sampling), not finding an infected deer in a sample of ∼1189 harvested deer from Franklin County would translate to an upper limit of CWD prevalence of > 5% (> 500 infected deer).

Importantly, our model-based framework supports an adaptive management approach. Surveillance strategies for target populations can be iteratively improved by updating the models with current information on host demography, harvest patterns and disease prevalence. Furthermore, the model landscape can readily incorporate locale-specific geographic features to better simulate host, harvest and disease distribution. Using the example of CWD surveillance in white-tailed deer of Missouri, we have illustrated how the model-based framework can guide collection and analysis of surveillance data that relies on non-probabilistic methods like harvest-based sampling. Although the models were designed with the needs of one particular agency in mind, they can easily be modified to be used by other agencies dealing with similar issues. The broader framework can be adapted for surveillance of CWD in white-tailed deer populations of other regions, or for surveillance of other important diseases of white-tailed deer like bovine tuberculosis and epizootic hemorrhagic disease. The framework can also be adapted to other host-pathogen systems and used for informed-decision making while planning and implementing disease surveillance in wild and free-ranging species.

## Acknowledgments

We thank and acknowledge D. Corcoran Barrios, J. Batten, T. Bixler, E. Flinn, V. Grimm, J. McRoberts, C. Miller and C. Scroggins for useful suggestions. Support for this study came from the Missouri Department of Conservation. AB was also partly supported by NSF Award P20GM104420.

## References

1. Drew L, Jessup A, Burr A, Franti CE. Serologic survey for brucellosis in feral swine, wild ruminants, and black bear of California, 1977 to 1989. J Wildl Dis. 1992;28: 355–363. doi:10.7589/0090-3558-28.3.355

2. Muñoz PM, Boadella M, Arnal M, de Miguel MJ, Revilla M, Martínez D, et al. Spatial distribution and risk factors of brucellosis in Iberian wild ungulates. BMC Infect Dis. 2010;10: 46. doi:10.1186/1471-2334-10-46

3. Dusek RJ, Bortner JB, DeLiberto TJ, Hoskins J, Franson JC, Bales BD, et al. Surveillance for high pathogenicity avian influenza virus in wild birds in the Pacific Flyway of the United States, 2006-2007. Avian Dis. 2009;53: 222–230. doi:10.1637/8854.1

4. Dusek RJ, Hagen CA, Franson JC, Budeau DA, Hofmeister EK. Utilizing hunter harvest effort to survey for wildlife disease: A case study of West Nile virus in greater sage-grouse. Wildl Soc Bull. 2014;38: 721–727. doi:10.1002/wsb.472

5. Samuel MD, Joly DO, Wild MA, Wright SD, Otis DL, Werge RW, et al. Surveillance strategies for detecting chronic wasting disease in free-ranging deer and elk: results of a CWD surveillance workshop, 10-12 December 2002. United States Geol Surv Natl Wildl Heal Center, Madison, Wisconsin, USA. Madison, Wisconsin, USA; 2003; 41.

6. Rees EE, Merrill EH, Bollinger TK, Hwang Y Ten, Pybus MJ, Coltman DW. Targeting the detection of chronic wasting disease using the hunter harvest during early phases of an outbreak in Saskatchewan, Canada. Prev Vet Med. Elsevier B.V.; 2012;104: 149–159. doi:10.1016/j.prevetmed.2011.10.016

7. Haley NJ, Hoover EA. Chronic wasting disease of cervids: Current knowledge and future perspectives. Annu Rev Anim Biosci. 2015;3: 305–325. doi:10.1146/annurev-animal-022114-111001

8. Samuel MD, Storm DJ. Chronic wasting disease in white-tailed deer: infection, mortality and implications for heterogeneous transmission. Ecology. 2016;97: 3195–3205.

9. Gross JE, Miller MW. Chronic wasting disease in mule deer: disease dynamics and control. J Wildl Manage. 2001;65: 205–215. doi:10.2307/3802899

10. Edmunds DR, Kauffman MJ, Schumaker BA, Lindzey FG, Cook WE, Kreeger TJ, et al. Chronic wasting disease drives population decline of white-tailed deer. PLoS One. 2016;11: e0161127. doi:10.1371/journal.pone.0161127

11. Cannon RM, Roe RT. Livestock disease surveys. A field manual for veterinarians. Bureau of Resource Science, Department of Primary Industry. Aust. Govt. Publ. Serv. Canberra; 1982.

12. Fosgate GT. Practical sample size calculations for surveillance and diagnostic interventions. J Vet Diagnostic Investig. 2009;21: 3–14. doi:10.1177/104063870902100102

13. Nusser SM, Clark WR, Otis DL, Huang L. Sampling considerations for disease surveillance in wildlife populations. J Wildl Manage. 2008;72: 52–60. doi:10.2193/2007-317

14. Beringer J, Hansen L, Millspaugh J, Meyer T. A statewide surveillance effort for detecting chronic wasting disease in wild white-tailed deer in Missouri. Wildl Soc Bull. 2003;31: 873–881. Available: http://www.jstor.org/stable/3784613

15. ProMED-Mail. Chronic wasting disease, cervid - USA (02):(MISSOURI). ProMed-mail 20120125.1020977. 2012.

16. ProMED-Mail. Chronic wasting disease, cervid - USA (22): (Missouri). ProMed-mail 20151217.3870576. 2015.

17. CWD surveillance and monitoring in Missouri [Internet]. [cited 21 Dec 2017]. Available: https://huntfish.mdc.mo.gov/hunting-trapping/wildlife-diseases/chronic-wasting-disease-cwd/cwd-surveillance-and-monitoring

18. Walsh DP. Enhanced surveillance strategies for detecting and monitoring chronic wasting disease in free-ranging cervids: [Internet]. U.S. Geological Survey Open-File Report. 2012. Available: http://pubs.usgs.gov/of/2012/1036/

19. Joly DO, Samuel MD, Langenberg JA, Blanchong JA, Batha CA, Rolley RE, et al. Spatial epidemiology of chronic wasting disease in Wisconsin white-tailed deer. J Wildl Dis. 2006;42: 578–588. doi:10.7589/0090-3558-42.3.578

20. Osnas EE, Heisey DM, Rolley RE, Samuel MD. Spatial and temporal patterns of chronic wasting disease: Fine-scale mapping of a wildlife epidemic in Wisconsin. Ecol Appl. 2009;19: 1311–1322. doi:10.1890/08-0578.1

21. VerCauteren KC, Hygnstrom SE. White-tailed deer. In: Wishart D, editor. Encyclopedia of the Great Plains. Lincoln, NE: University of Nebraska Press; 2004. pp. 642–643.

22. Walter WD, VerCauteren KC, Campa H, Clark WR, Fischer JW, Hygnstrom SE, et al. Regional assessment on influence of landscape configuration and connectivity on range size of white-tailed deer. Landsc Ecol. 2010;24: 1405–1420. doi:10.1007/s10980-009-9374-4

23. Ozoga J. Whitetail intrigue. Krause, Iola, Wisconsin USA; 2000.

24. Monteith KL, Sexton CL, Jenks JA, Terry Bowyer R. Evaluation of techniques for categorizing group membership of white-tailed deer. J Wildl Manage. 2007;71: 1712–1716.

25. Ozoga J., Verme L., Bienz C. Parturition behavior and territoriality in white-tailed deer: Impacts on neonatal mortality. J Wildl Manage. 1982;46: 1–11.

26. Schwede G, Hendrichs H, McShea W. Social and spatial organization of female white-tailed deer, Odocoileus virginianus, during the fawning period. Anim Behav. 1993;45: 1007–1017.

27. Hirth D. Social behavor of white-tailed deer in relation to habitat. Wildl Monogr. 1977;53: 1–55.

28. Cullingham CI, Merrill EH, Pybus MJ, Bollinger TK, Wilson GA, Coltman DW. Broad and fine-scale genetic analysis of white-tailed deer populations: Estimating the relative risk of chronic wasting disease spread. Evol Appl. 2011;4: 116–131. doi:10.1111/j.1752-4571.2010.00142.x

29. Nelson ME. Natal dispersal and gene flow in white-tailed deer in northeastern Minnesota. J Mammal. 1993;74: 316–322.

30. Purdue JR, Smith MH, Patton JC. Female philopatry and extreme spatial genetic heterogeneity in white-tailed deer. J Mammal. 2000;81: 179–185.

31. Lutz CL, Diefenbach DR, Rosenberry CS. Population density influences dispersal in female white-tailed deer. J Mammal. 2015;96: 494–501.

32. Long ES, Diefenbach DR, Rosenberry CS, Wallingford BD. Multiple proximate and ultimate causes of natal dispersal in white-tailed deer. Behav Ecol. 2008;19: 1235–1242.

33. Lutz CL, Diefenbach DR, Rosenberry CS. Proximate influences on female dispersal in white-tailed deer. J Wildl Manage. 2016;80: 1218–1226.

34. Wilensky U. NetLogo [Internet]. Center for Connected Learning and Computer-Based Modeling, Northwestern University, Evanston, IL; 1999. Available: http://ccl.northwestern.edu/netlogo/.

35. Grimm V, Berger U, Bastiansen F, Eliassen S, Ginot V, Giske J, et al. A standard protocol for describing individual-based and agent-based models. Ecol Modell. 2006;198: 115–126.

36. Grimm V, Berger U, DeAngelis DL, Polhill JG, Giske J, Railsback SF. The ODD protocol: A review and first update. Ecol Modell. 2010;221: 2760–2768. doi:10.1016/j.ecolmodel.2010.08.019

37. Belsare A V., Gompper ME, Millspaugh JJ. MOOvPOP (Version 8). In: CoMSES Computational Model Library [Internet]. 2017. doi:https://doi.org/10.25937/cnex-s628

38. Belsare A V., Gompper ME, Millspaugh JJ. MOOvPOPsurveillance. In: CoMSES Computational Model Library [Internet]. 2017. doi:https://doi.org/10.25937/8hpz-9y96

39. Diefenbach DR, Long ES, Rosenberry CS, Wallingford BD, Smith DR. Modeling distribution of dispersal distances in male white-tailed deer. J Wildl Manage. 2008;72: 1296–1303. doi:http://dx.doi.org/10.2193/2007-436

40. Long ES, Diefenbach DR, Rosenberry CS, Wallingford BD, Grund MD. Forest cover influences dispersal distance of white-tailed deer. J Mammal. 2005;86: 623–629. doi:http://dx.doi.org/10.1644/1545-1542(2005)86[623:FCIDDO]2.0.CO;2

41. Ditchkoff SS. Anatomy and Physiology. In: Hewitt DG, editor. Biology and management of white-tailed deer. Boca Raton, FL: CRC Press; 2011. pp. 43–73.

42. Hiller TL, Henry Campa I, Winterstein SR. Survival and space use of fawn white-tailed deer in southern Michigan. Am Midl Nat. 2008;159: 403–412.

43. Van Deelen TR, Campa Iii H, Haufler JB, Thompson PD. Mortality patterns of white-tailed deer in Michigan’s Upper Peninsula. J Wildl Manage. 1997;61: 903–910.

44. Marchinton RL, Hirth D. Behavior. In: Halls LK, editor. White-tailed deer: Ecology and management. Harrisburg, PA: Stackpole Books; 1984. pp. 129–168.

45. VerCauteren KC, Hygnstrom SE. Managing White-tailed Deer: Midwest North America. In: Hewitt DG, editor. Biology and management of white-tailed deer. 1st ed. Boca Raton, FL: CRC Press; 2011. pp. 501–535.

46. Nelson ME, Mech LD. Dispersal in female white-tailed deer. J Mammal. 1992;73: 891–894.

47. Miller BF, DeYoung R., Campbell TA, Laseter BR, Ford WM, Miller KV. Fine-scale genetic and social structuring in a central Appalachian white-tailed deer herd. J Mammal. 2010;93: 681–689.

48. Smith, W P. Odocoileus virginianus. Mamm Species. 1991;388: 1–13.

49. Miller KV, Muller LI, Demarias S. White-tailed deer (Odocoileus virginianus). In: Feldhamer GA, Thompson BC, Chapman JA, editors. Wild mammals of North America: Biology, management, and conservation. 2nd ed. Baltimore, Maryland: Johns Hopkins University Press; 2003. pp. 906–930.

50. Gaillard JM, Festa-Blanchet M, Yoccoz N., Loison A, Toigo C. Temporal variation in fitness components and population dynamics of large herbivores. Annu Rev Ecol Syst. 2000;31: 367–393.

51. Nixon CM, Mankin PC, Etter DR, Hansen LP, Brewer PA, Chelsvig JE, et al. White-tailed deer dispersal behavior in an agricultural environment. Am Midl Nat. 2007;157: 212–220.

52. Bevins SN, Dusek RJ, White CL, Gidlewski T, Bodenstein B, Mansfield KG, et al. Widespread detection of highly pathogenic H5 influenza viruses in wild birds from the Pacific Flyway of the United States. Sci Rep. 2016;6: 28980. doi:10.1038/srep28980

53. Schöning JM, Cerny N, Prohaska S, Wittenbrink MM, Smith NH, Bloemberg G, et al. Surveillance of bovine tuberculosis and risk estimation of a future reservoir formation in wildlife in Switzerland and Liechtenstein. Gordon S V., editor. PLoS One. 2013;8: e54253. doi:10.1371/journal.pone.0054253

54. O’Brien DJ, Schmitt SM, Berry DE, Fitzgerald SD, Lyon TJ, Vanneste JR, et al. Estimating the true prevalence of Mycobacterium bovis in free-ranging elk in Michigan. J Wildl Dis. 2008;44: 802–810.

55. Rees EE, Merrill EH, Bollinger TK, Hwang Y Ten, Pybus MJ, Coltman DW. Targeting the detection of chronic wasting disease using the hunter harvest during early phases of an outbreak in Saskatchewan, Canada. Prev Vet Med. Elsevier B.V.; 2012;104: 149–159. doi:10.1016/j.prevetmed.2011.10.016

56. Diefenbach DR, Rosenberry CS, Boyd RC. Efficacy of detecting chronic wasting disease via sampling hunter-killed white-tailed deer. Wildl Soc Bull. 2004;32: 267–272.

57. Schmitt SM, Fitzgerald SD, Cooley TM, Bruning-Fann CS, Sullivan L, Berry D, et al. Bovine tuberculosis in free-ranging white-tailed deer from Michigan. J Wildl Dis. 1997;33: 749–758.

58. Bevins SN, Dusek RJ, White CL, Gidlewski T, Bodenstein B, Mansfield KG, et al. Widespread detection of highly pathogenic H5 influenza viruses in wild birds from the Pacific Flyway of the United States. Sci Rep. 2016;6: 28980. doi:10.1038/srep28980

