## Supplementary figures and images for "An agent-based framework for improving wildlife disease surveillance: A case study of chronic wasting disease in Missouri white-tailed deer"

### Figure 9

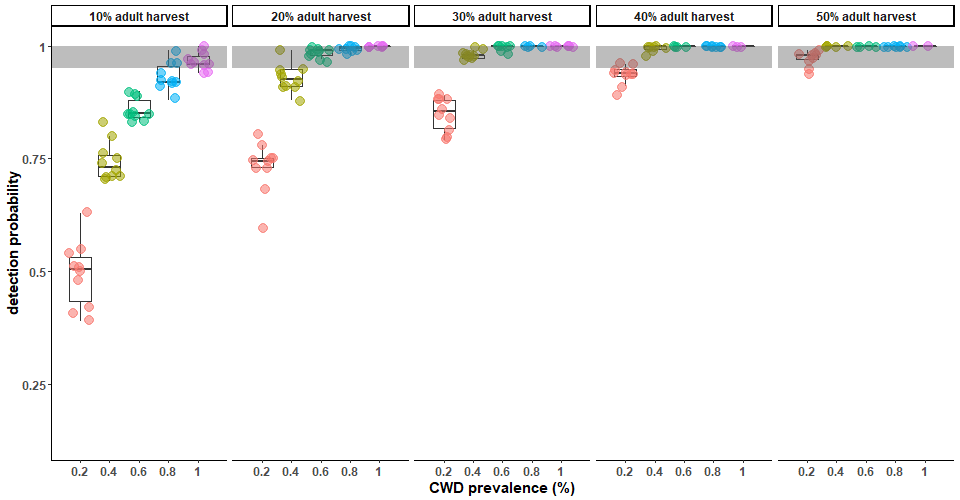
